# Multiple roles for *laccase2* in butterfly wing pigmentation, scale development, and cuticle tanning

**DOI:** 10.1101/858167

**Authors:** Ceili L. Peng, Anyi Mazo-Vargas, Benjamin J. Brack, Robert D. Reed

**Affiliations:** Department of Ecology and Evolutionary Biology, Cornell University, Ithaca, NY 14850

**Keywords:** melanin, *Vanessa cardui*, Lepidoptera, CRISPR, pigmentation, color pattern

## Abstract

Lepidopteran wing scales play important roles in a number of functions including color patterning and thermoregulation. Despite the importance of wing scales, however, we still have a limited understanding of the genetic mechanisms that underlie scale patterning, development, and coloration. Here we explore the function of the phenoloxidase-encoding gene *laccase2* in wing and scale development in the nymphalid butterfly *Vanessa cardui*. Somatic deletion mosaics of *laccase2* generated by CRISPR/Cas9 genome editing presented several distinct mutant phenotypes. Consistent with work in other non-lepidopteran insect groups, we observed reductions in melanin pigmentation and defects in cuticle formation. We were also surprised, however, to see distinct effects on scale development including complete loss of wing scales. This work highlights *laccase2* as a gene that plays multiple roles in wing and scale development and provides new insight into the evolution of lepidopteran wing coloration.

## 1 INTRODUCTION

Wing scales are a defining feature of the insect order Lepidoptera. These complex and diverse structures play several important physiological and ecological adaptive functions in the life histories of moths and butterflies, and are the color bearing units of wing color patterns themselves. Although progress has recently been made in understanding the genetic basis of color patterning in butterfly wings (e.g., Mazo-Vargas et al., 2017; Zhang et al., 2017a,b; Lewis et al., 2019), we still have a limited understanding of the genetics of color pigmentation itself. In the last few years, some progress has been made in functionally characterizing wing scale pigmentation genes using CRISPR/Cas9 genome editing (Li et al., 2015; Zhang et al., 2017a; Matsuoka & Monteiro, 2018.). Despite this recent work, however, there are still many pigmentation genes that need to be characterized if we are to gain a more thorough understanding of the origin and evolution of wing coloration. One of these genes is *laccase2* (*lac2*), which is the focus of this study.

*lac2* encodes a phenoloxidase that oxidizes catecholenines into quinones (Hopkins & Kramer, 1992). RNAi-mediated *lac2* knockdowns have demonstrated that this gene plays a role in both cuticle formation and pigmentation in a range of insects. Stinkbugs (Futahashi et al., 2011), milkweed bugs (Liu et al., 2014), honey bees (Elias-Neto et al., 2010), mosquitos (Du et al., 2017), pine sawyer beetles (Niu et al., 2008), and red flour beetles (Arakane et al., 2005) that experienced *lac2* knockdown were unable to properly develop pigmented cuticle, resulting in soft, demelanized bodies. *Drosophila melanogaster* affected by *lac2* knockdown fail to melanize and also improperly develop wing cuticle and wing sensory bristles (Riedel et al., 2011). *lac2* knockdown in mosquitoes also results in eggs that are pale and less durable, resulting in high mortality (Wu et al., 2013). Thus, there is extensive evidence for a deeply conserved role for *lac2* in insect cuticle development.

To our knowledge *lac2* remains poorly characterized characterized in Lepidoptera – likely due to the technical difficulties of RNAi in this group (Terenius et al., 2011). Nonetheless, expression studies have suggested that *lac2* may play some roles in lepidopteran development. There is strong expression of *lac2* during cuticle sclerotization developmental stages in tobacco hornworms (Dittmer et al., 2004). Also, *lac2* expression is associated with some melanin larval color markings in *Papilio xuthus* and *Bombyx mori* (Futahashi, Banno, & Fujiwara, 2010). Outside of larval color associations in these species, however, potential pigmentation-related roles of *lac2* in Lepidoptera are not well understood.

Here we use CRISPR/Cas9 somatic mosaic knockouts to functionally characterize the role *lac2* plays during wing development of the butterfly *Vanessa cardui*. We observed a variety of phenotypic effects ranging from compromised pigmentation to complete loss of scales. This work highlights *lac2* as an important gene that plays multiple functions in butterfly wing and color pattern development, and helps expand our current models of butterfly wing pattern evolution and development.

## 2 MATERIALS AND METHODS

### 2.1 Butterfly culture

Our *V. cardui* butterfly colony originated from Carolina Biological Supply. Butterflies were maintained in a growth chamber (at 27°C and 16 hours of daylight) and fed 10% sugar water. *V. cardui* were induced to oviposit on fresh-cut *Malva parviflora* leaves. The larvae were fed a combination of *M. parviflora* and artificial diet supplied by Carolina Biological Supply.

### 2.2 CRISPR/Cas9 genome editing

In order to test of the function of *lac2* in butterfly wing development we followed the standard CRISPR/Cas9 G_0_ somatic deletion mosaic approach used in most previous butterfly wing CRISPR studies. Briefly, syncytial embryos are injected with sgRNA and Cas9, resulting in individuals genetically mosaic for Cas9-mediated deletions. A subset of these injected individuals survive to adulthood and display mutant clones (i.e., clusters of cells bearing the induced mutation). Cas9-induced mutant clones can be unambiguously identified because they are asymmetrical between left and right sides of the butterfly. Natural variation and family effects (e.g., germ line mutations) present symmetrical phenotypes, and asymmetrical mutant phenotypes are never observed in water or Cas9-only negative control injections. For this study we followed the specific protocol of Zhang and Reed (2017). Single guide RNAs (sgRNAs) were designed using the template of GGN_18_NGG. For this experiment two sgRNAs were used to facilitate long deletions. The sequences of each of the sgRNAs are as follows, with complementary cut site sequences underlined in bold:

sgRNA 1: 5’ – GAAATTAATACGACTCACTATA<u>**GGGAACAGCGACGCCTTTCC**</u>GTT TTAGAGCTAGAAATAGC
sgRNA 2: 5’-GAAATTAATACGACTCACTATA<u>**GGCGGGGCAAACTGAAGCAA**</u>GTT TTAGAGCTAGAAATAGC

sgRNAs were designed to have a high GC content (>60%), to be no more than 2 kb apart, and to have a range over multiple exons if possible. In this case, the *lac2* gene consists of 8 exons spread across 48,895 bp. The sgRNAs were designed to cleave the DNA at 39,747 bp and 41,662 bp, to create a 1915 bp deletion that starts in exon 2, completely removes exon 3, and ends in exon 4 (Fig. 1a). The sgRNA template was amplified by PCR using a forward primer that incorporated a T7 polymerase-binding site and a sgRNA target site, and a reverse primer that encoded the sgRNA sequence. The MegaScript T7 kit (Ambion) was then used to perform *in vitro-*transcription. The sgRNAs were purified using phenol/chloroform extraction and isopropanol precipitation.

**Figure 1.**
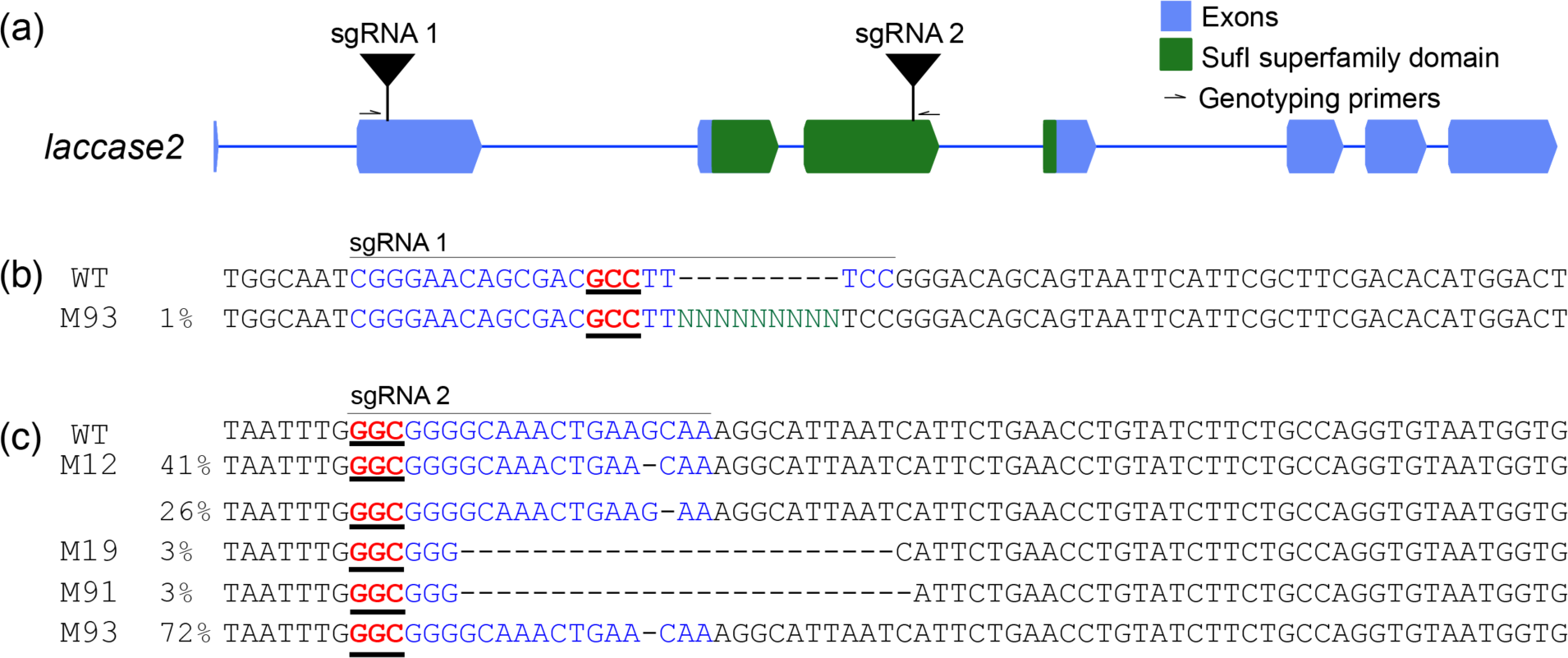
CRISPR editing strategy for *lac2*. (a) The *V. cardui lac2* locus, annotated with sgRNA and genotyping primer sites as described in the text. Genotyping of mutant individuals confirmed lesions at (b) sgRNA1 and (c) sgRNA2 sites. Individual mutant names (M#) and ICE-estimated mutant allele frequencies are reported to the left of each sequence.

Prior to injection, 0.5 µl of 1000 ng/µl Cas9 protein (PNABio, catalog number CP01) was combined with both sgRNAs to create a solution with a final concentration of 330 - 480 ng/µl of each sgRNA and 333 ng/µl of Cas9. To collect eggs, fresh *M. parvifola* leaves were placed in a *V. cardui* colony cage for 1-3 h to induce oviposition. The resulting eggs were then lined up on strips of double-sided adhesive tape on slides and exposed to desiccant for 20 minutes. Needles pulled from borosilicate glass capillaries (Sutter Instrument Company, 0.5 mm, catalog number BF-100-50-10) were used for injection. Egg injections were performed between 2-4 h of oviposition. Injected eggs were incubated in a growth chamber at 27°C until hatching. Once hatched, larvae were transferred to either diet, cut *M. parviflora* leaves, or live plants, and raised in small groups.

### 2.3 Mutant genotyping

To confirm deletions, two primers were generated to flank the targeted *lac2* deletion interval (Fig. 1):

Primer 1: 5’-TCGTGACAACATTCGCTGGA
Primer 2: 5’-AACTTTGACTGGTTGCACCG

Genomic DNA was extracted from adult butterfly thoracic muscles using proteinase K as a digestion buffer. As in previous butterfly wing-focused studies, muscle tissue was used for genotyping since wing scales of adult butterflies are dead cuticle and no longer have sufficient intact DNA for sequence characterization of mutant alleles (see Zhang and Reed, 2017). The primers were used to amplify the Cas9 target region via PCR, and the size of the fragments was detected using gel electrophoresis. The genotyping primers generate a product 2287 bp long fragment in a wild type genome, and it is expected that the amplified mutated gene sequence will be ∼357 bp long if the expected long deletion and repair occurred. To identify short insertions or deletions produced by a single guide RNA, DNA fragments were isolated from wild type and mutant butterflies and were extracted from an agarose gel. These fragments were purified and sequenced with Sanger sequencing using the primers above. Sequences from mutant butterflies were then characterized using the Inference of CRISPR Edits (ICE) tool (Hsiau et al., 2018) to identify and quantify mutant alleles.

## 3 RESULTS

We injected 418 *V. cardui* eggs with Cas9 and sgRNAs. Of those eggs, 145 larvae hatched resulting in a hatch rate of 35%. 72 butterflies were ultimately raised to adulthood and assessed for mosaic phenotypes. Wings were examined for mutations that appeared clonal, meaning that sections of wings were uniformly different and showed distinct boundaries compared to surrounding areas, and asymmetrical (i.e. left/right wings were different from each other). We identified 5 butterflies with distinct melanin-deficient wing clones, and 9 with scaleless wing clones. We also noted 10 butterflies with deformed wings. The eyes and cuticle of adult butterflies were also inspected for obvious mutations and none were observed in these experiments, although we caution that these negative results do not necessarily rule out a role for *lac2* in adult non-wing cuticle development, especially given the mosaic nature of our assay. To confirm accurate targeting of sgRNAs we sequenced *lac2* from mutant butterflies and positively identified alleles with lesions at the predicted sites (Fig. 1b,c), and confirmed multiple long deletions by gel electrophoresis. Overall, we observed three different categories of phenotypes: pigmentation loss (reduction of pigmentation in scales, loss of larval pigmentation), missing scales, and deformed wings.

The first major category of *lac2* mosaic mutants had clonal patches of reduced pigmentation (n=5) (Fig. 2a). This effect was most frequently observed in black pattern elements of dorsal forewings, where melanic scale cells showed reduced pigment intensity (Fig. 2a, mutant M43). We also observed one example with a ventral hindwing phenotype, where a field of normally brown scales of uncertain pigment composition, exhibited a lighter beige coloration (Fig. 2a,b, mutant M60). In addition to effects on wing scales, we also noticed loss of melanin pigmentation in clonal patches of the larval head capsule (Fig. 2c), the larval soft integument, and cuticular spines, which were often malformed (n=2) (Fig. 2d).

**Figure 2.**
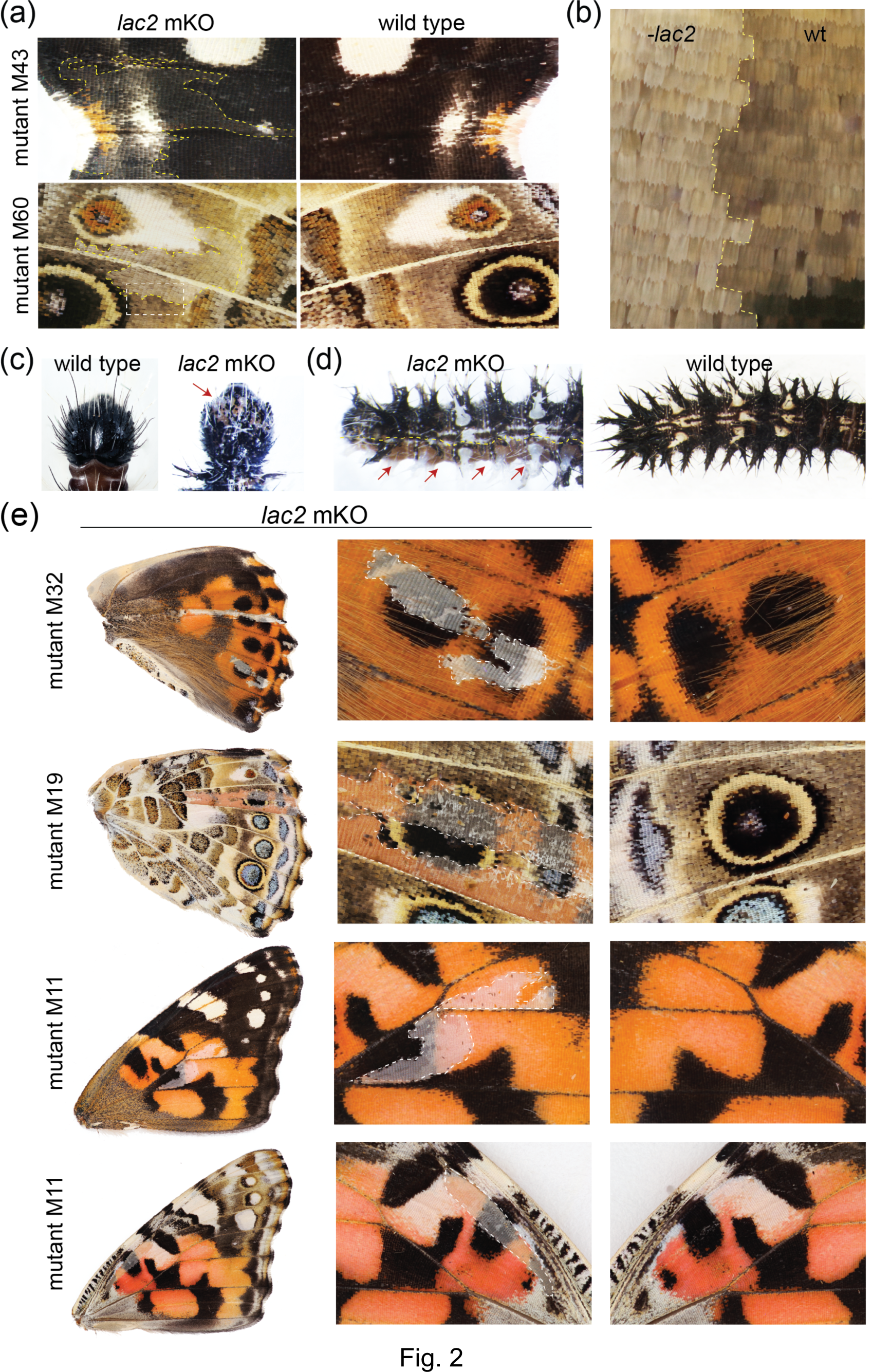
*lac2* mosaic knockouts (mKOs) result in pigmentation defects and scale loss. Wing mutants are asymmetric mosaics, with left-right comparisons from the same individual shown. (a) Wing scales show reduced pigmentation in both black and brown color patterns. Presumptive *lac2* deletion clone boundaries are annotated by dashed yellow lines. (b) High magnification image mutant M60, from white dashed box in panel (a), shows a boundary (yellow dashed line) between presumptive *lac2* deletion scales and wild type scales. (c) A larva with pigment-deficient clones in the head capsule cuticle, as highlighted by the red arrow. (d) A larva showing mKO pigment deficiency the larval integument along the bottom edge, as marked by the red arrows. (e) mKOs showing presumptive *lac2* deletion clones with loss of wing scales. These clones often cross pigmentation and color pattern boundaries and do not have obvious major effects on wing structure. Dashed lines mark presumptive clone boundaries.

Another common category of mutants were individuals with large swaths of missing wing scales (n=9) (Fig. 2e). These mutations were found on both dorsal and ventral sides of forewings and hindwings, and crossed pigment boundaries (e.g. melanin/ommochrome boundaries) and pattern boundaries, including through eyespots (e.g. Fig. 2e, M19). While scale loss can be caused by mechanical damage, in this case we are confident that scaleless clones were due to genetic mutation for several reasons. First, butterflies were frozen immediately after emergence to preclude wear or collision damage. Second, the boundary lines between scaled and scaleless patches are clean and precise, consistent with cell lineage (clonal) effects as seen in gynandromorphs, pigment gene knockout clones, etc. This is not consistent with scratches, wing deformities, or pupal emergence problems, which display ragged edges and many dislodged scales. Third, many of the mutant clones follow wing vein boundaries (e.g., all mutants in Fig. 2e), as is commonly seen in clonal mosaics in both butterfly and *Drosophila* wings – this would not be the case with mechanical damage where large scratches should cross wing veins. We speculate that the observed scale loss phenotypes are likely due of underlying cuticular structures that have been weakened by *lac2* knockout.

In addition to localized effects on pigmentation and scale development, we also observed that many presumptive mutant butterflies were unable to pupate correctly, and either failed to emerge, or emerged with crumpled wings (n=10). In some cases, we observed that the creases in these crumpled wings corresponded to clones with missing scales, suggesting an association between *lac2* knockout and structural robustness of the wing.

## 4 DISCUSSION

Here we analyzed the function of *lac2* in butterfly wing development using CRISPR/Cas9 mosaic knockouts. The clearest mutant phenotypes we observed manifested as color and structural defects to cuticular features, including wings, wing scales, the larval head capsule, and larval spines. Previous research shows that melanin pigmentation and cuticularization are often linked through a process called tanning, and can share a common set of genes and gene products (Hopkins & Kramer, 1992; Arakane et al., 2005; Du et al., 2017; Moussian, 2010). The cuticle is made up of several layers which follow an ordered synthesis. Once the cuticle has formed, a variety of compounds derived from tyrosine are released into the newly secreted cuticle. These compounds are then oxidized to form quinones which can be used for melanin pigmentation or for cuticular protein crosslinking and sclerotization (Riedel et al., 2011). Both of these functions – cuticle synthesis and pigmentation – depend on *lac2*, thus providing a potential explanation for our observed phenotypes that include both compromised cuticular structures (e.g., crumpled wings and deformed larval spines) as well as very specific effects on pigmentation (e.g., miscolored scales). This connection between cuticularization and melanization could also explain the dual effects of several other genes on both coloration and scale structure in butterflies, including *tyrosine hydroxylase, dopa decarboxylase, yellow, ebony*, and *aaNAT* (Zhang et al., 2017; Matsuoka & Monteiro, 2018). Thus, our *lac2* results expand and reinforce our emerging view of the close functional and genetic relationship between structure and color in insects.

One class of mutant phenotype we were surprised by was the specific and distinctive loss of scale cells. It is plausible that this could be related with strong, but highly localized, cuticularization defects that did not affect other wing features. There is precedent from *Drosophila*, however, of *lac2* mutants displaying a specific loss of wing bristles (Riedel et al., 2011). It is thought that lepidopteran wing scales were evolutionarily derived from ancestral sensory bristles (Galant et al., 1998), therefore there may be some special function of *lac2* in development of this general cell type that we do not fully understand. Whatever the case may be, the scale loss effects of *lac2* deletions only further highlights the central role this gene plays in many aspects of wing scale development. This new understanding of *lac2* function fills an important gap in our current models of the evolution and development of butterfly wing patterns.

## ACKNOWLEDGEMENTS

This work represents the undergraduate thesis research of Ceili Peng, and we would like to thank the Cornell EEB honors thesis program participants for support and feedback on the project. We would also like to thank two anonymous reviewers for helpful suggestions that improved the manuscript. This work was supported by U.S. National Science Foundation grants IOS-1557443 and IOS-1656514 to RDR.

